# Inflexible center-frequency shifts in neural speech tracking are linked to reading and phonological deficits in developmental dyslexia

**DOI:** 10.1101/2025.09.02.673698

**Authors:** Doris Gahleitner, Anne Hauswald, Nathan Weisz, Fabian Schmidt

**Affiliations:** Paris-Lodron-University of Salzburg, Department of Psychology, Centre for Cognitive Neuroscience, Salzburg, Austria; Neuroscience Institute, Christian Doppler University Hospital, Paracelsus Medical University, Salzburg, Austria

**Keywords:** developmental dyslexia, temporal sampling framework, neural speech tracking, speech-brain coherence, magnetoencephalography, spectral parameterization

## Abstract

Developmental dyslexia is marked by persistent deficits in reading and phonological awareness, potentially linked to atypical neural tracking of acoustic speech rhythms at frequencies below 10 Hz. Using magnetoencephalography, we compared adults with developmental dyslexia and matched controls while they listened to speech with varying levels of intelligibility (manipulated via noise vocoding). In controls, decreasing intelligibility led to systematic increases in the tracked frequency of the speech stream, replicating previous findings. This increase in tracking frequency was largely absent in individuals with developmental dyslexia. Unexpectedly, the magnitude of periodic speech tracking in the low frequency range was preserved in DD. Instead, the increase in central tracking frequency was linked to phonological awareness and reading performance.

## 1. Introduction

Reading and writing are important skills for educational success, career prospects and overall well-being (Hunter & Leiper, 1993; Zajacova & Lawrence, 2018). Difficulties in acquiring these skills can negatively affect academic achievement, self-concept and mental health (Hendren et al., 2018; Wilmot, Hasking, et al., 2023; Wilmot, Pizzey, et al., 2023). Developmental dyslexia (DD) is the most prevalent specific learning disorder, affecting about 7% of schoolchildren worldwide (American Psychiatric Association, 2013; Yang et al., 2022). It is characterised by substantial and persistent reading difficulties not attributable to a more general intellectual impairment, visual acuity problems or inadequate schooling (American Psychiatric Association, 2013; World Health Organization, 2022).

Core predictors of DD include verbal short-term memory, rapid automatised naming and, most prominently phonological awareness (Habib & Giraud, 2013; Ptok et al., 2008; Caravolas et al., 2012; Landerl et al., 2019; Melby-Lervåg, 2012; Vibulpatanavong & Evans, 2019) – the ability to identify and manipulate speech segments such as syllables or phonemes (Anthony & Francis, 2005; Schnitzler, 2015). Phonological awareness is particularly crucial for early reading acquisition (Melby-Lervåg, 2012; Ptok et al., 2008; Schnitzler, 2015) and remains impaired in individuals with DD well into their adulthood (Bruck, 1992; Peterson & Pennington, 2012; Reis et al., 2020). The persistence of this deficit points to a deeper disruption in the neural processing of speech in individuals with DD.

The Temporal Sampling Framework (TSF, Goswami, 2011) provides one influential account of this disruption. It suggests that DD arises from reduced perceptual sensitivity to slow rhythmic acoustic amplitude modulations (<10 Hz) in speech. These modulations provide prosodic (∼0.5-3 Hz, delta) and syllabic (∼4-8 Hz; theta) cues for segmenting a “continuous” acoustic signal into linguistically meaningful units (Goswami, 2011, 2018, 2019; Poeppel & Assaneo, 2020). The TSF therefore predicts that this reduced sensitivity should be observable neurally as a decrease in coherence or phase-locking to the speech envelope in DD (Abrams et al., 2009; Molinaro et al., 2016; Goswami, 2011, 2018). During brain maturation, such reduced alignment of neural oscillations in these frequency bands to acoustic cues may compromise the formation of stable phonological representations. Behavioral evidence for this view includes impaired synchronization to metronome beats (Thomson & Goswami, 2008), difficulties in perceiving musical beats, amplitude rise times, syllabic stress (Kalashnikova et al., 2021, Huss et al., 2011, Law et al., 2014; Leong et al., 2011; Thomson & Goswami, 2010; Van Hirtum et al., 2019) and a reduced speech in noise perception (Van Hirtum et al., 2021; Ziegler et al., 2009). Together, these findings point toward a sensory-level impairment in neural processing of acoustic information in DD, which the TSF attributes to reduced sampling of rhythmicity in acoustic stimuli via oscillatory neural activity (Goswami, 2019).

Despite its appeal, the TSF faces several challenges. For one, speech is not strictly rhythmic (Voss & Clarke, 1975), while some rhythmicity can be observed in the theta range (related to the production of syllables; Coupé et al., 2019; Giraud & Poeppel, 2012; Gross et al., 2013; Hyafil et al., 2015; Meyer, 2018), delta band fluctuations in everyday continuous natural speech are not particularly rhythmic (Meyer et al., 2020). Nevertheless, reduced neural speech tracking in DD has more frequently been found in the delta than in the theta range (Araújo et al., 2024; Lizarazu et al., 2021; Zhang et al., 2021; 2022). Relatedly, most studies that test the TSF have relied on canonical frequency bands without distinguishing between the neural tracking of rhythmic and arrhythmic (i.e. aperiodic/broadband) components in acoustic signals. This risks misattributing broadband to narrowband rhythmic processes (Donoghue et al., 2020; Schmidt et al., 2023). Furthermore, neural tracking of speech is not necessarily a static process and the information being tracked may change depending on attentional focus or cognitive demands. For instance, speech intelligibility modulates neural tracking in systematic ways. When speech is clear, coherence peaks tend to align with the syllabic rate (∼4 Hz). Under degraded conditions—for example, when speech is noise-vocoded—these peaks shift toward faster envelope fluctuations (∼5–7 Hz; see Schmidt et al., 2023 & Chen et al. 2023). This suggests that the tracking of rhythmic fluctuations in speech is not simply reduced by decreasing signal quality, but that the brain flexibly prioritizes different temporal cues in the acoustic signal depending on how intelligible the speech is (Schmidt et al., 2023). Importantly, without parametrization, such increases in center frequency can appear as reductions in coherence magnitude. This raises the possibility that what has often been described as reduced neural tracking – in work building on the TSF – may in fact reflect a lack of flexibility in adapting neural sampling to changing acoustic cues (Goswami, 2011, 2018).

Building on these insights, we investigated whether individuals with DD differ from matched controls in their neural tracking of speech across varying levels of intelligibility, manipulated via noise vocoding (Chen et al., 2023; Hauswald et al., 2022; Schmidt et al., 2023). To obtain a more fine-grained view of neural-speech tracking, we parameterized speech-brain coherence spectra in their periodic (rhythmic) and aperiodic components (Schmidt et al., 2023; Chen et al., 2023). This approach enabled us to test whether DD is characterized by reduced flexibility in shifting between syllabic and envelope timescales, as well as by overall reductions in tracking strength, as predicted by the TSF. We further examined whether individual differences in such flexibility relate to reading and phonological skills. Our preregistered hypotheses are available at https://osf.io/9f2xg.

In controls, decreasing intelligibility produced systematic increases in the center frequency of coherence peaks, replicating previous findings (Schmidt et al. 2023, Chen et al. 2023). This adaptive shift was largely absent in the DD group. Moreover, shift magnitude reliably correlated with phonological awareness and reading performance, making it the most informative spectral feature we tested. Contrary to our TSF-based predictions, the overall strength of periodic low frequency neural tracking showed no significant group differences at any intelligibility level.

## 2. Results

### 2.1 Individuals with developmental dyslexia differ from controls on phonological and reading related abilities, but are matched in age, hearing and their vocabulary

In order to be able to interpret potential differences in neural tracking, it was important at the outset to evaluate how our two recruited groups of participants (DD and matched controls) differ on a DD related skillset. We therefore compared the two groups regarding reading skill and two self-developed measures for phonological awareness (Schnitzler, 2015; Stock et al., 2017) three possible confounding variables (Table 1). During word-reading and pseudoword-reading, subjects had to read as many words or pseudowords as they could within one minute (Moll & Landerl, 2014). In the self-developed measures for phonological awareness “spoonerism” and “syllable counting” tasks subjects were asked to solve 27 spoonerisms and count the syllables of 20 pseudowords (syllable counting) as quickly as possible (see *Methods – Cognitive Tasks* for details). For both “syllable counting” and “spoonerism’s” we calculated the inverse efficiency scores (IES), a measure for the “average energy consumed by the system over trials” (Townsend & Ashby, 1983). Across both the reading and the phonological awareness related tasks subjects with DD performed worse than controls (see Table 1). Additionally, our self-developed measures for phonological awareness were related to more established markers of reading performance, highlighting the link between phonological awareness and word reading performance: spearman’s rank-order correlations (*r*□) revealed negative relationships between word reading speed and the inverse efficiency scores on phonological awareness (*r*1 ranging from -0.61 on “spoonerism” to -0.52 on “syllable counting”, all *p*□<0.05). A similar observation was made for pseudoword reading: phonological awareness tasks were negatively related to pseudoword reading speed (*r*□ ranging from -0.50 on “spoonerism” to -0.60 on “syllable counting”; all *p*□<0.05). Neither age, hearing ability nor vocabulary showed a significant relationship with any of the cognitive tasks (all *p*1>0.2). Importantly, subjects with DD did not differ from controls based on their vocabulary (assessed via WAIS-D; *BF*_10_ = 0.34), age (*BF*_10_ = 0.39) or hearing ability (*BF*_10_ = 0.39; see also Table 1).

**Table 1.**
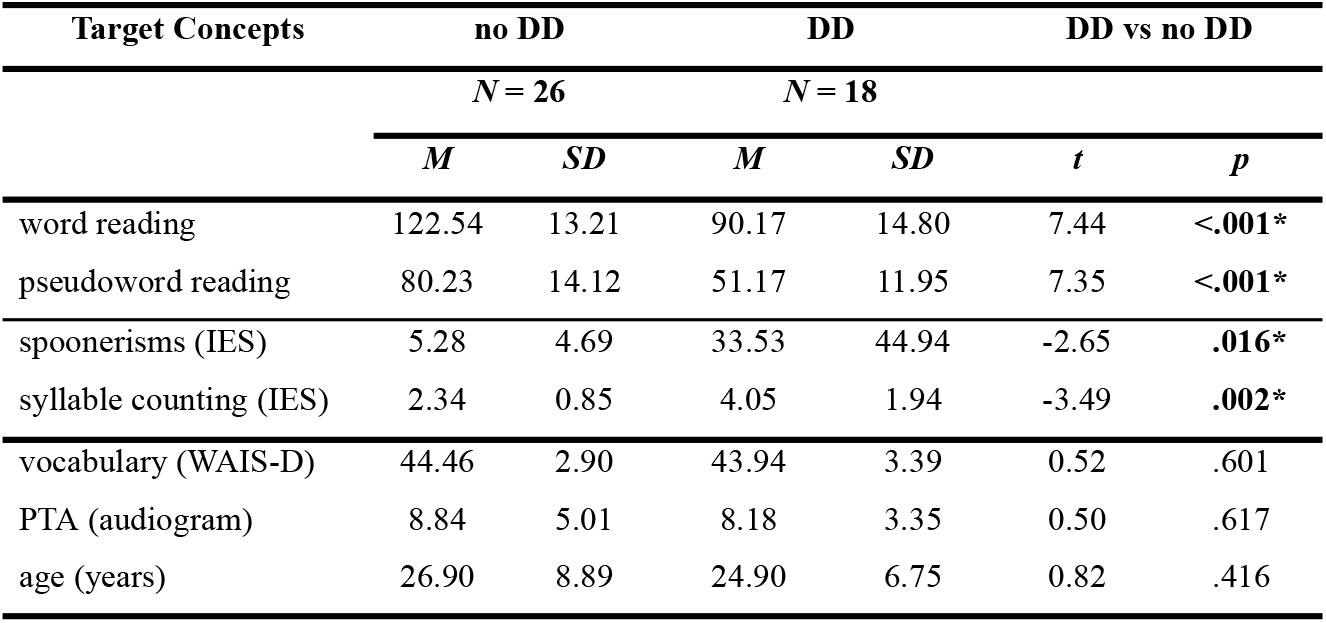
Descriptives on all Outcomes of the 4 Cognitive Tasks per DD Group, as well as control variables such as age, hearing ability (pure-tone average audiogram) and vocabulary.

| Target Concepts | no DD |  | DD |  | DD vs no DD |  |
| --- | --- | --- | --- | --- | --- | --- |
|  | *N* = 26 |  | *N* = 18 |  |  |  |
|  | *M* | *SD* | *M* | *SD* | *t* | *p* |
| word reading | 122.54 | 13.21 | 90.17 | 14.80 | 7.44 | <.001* |
| pseudoword reading | 80.23 | 14.12 | 51.17 | 11.95 | 7.35 | <.001* |
| spoonerisms (IES) | 5.28 | 4.69 | 33.53 | 44.94 | -2.65 | .016* |
| syllable counting (IES) | 2.34 | 0.85 | 4.05 | 1.94 | -3.49 | .002* |
| vocabulary (WAIS-D) | 44.46 | 2.90 | 43.94 | 3.39 | 0.52 | .601 |
| PTA (audiogram) | 8.84 | 5.01 | 8.18 | 3.35 | 0.50 | .617 |
| age (years) | 26.90 | 8.89 | 24.90 | 6.75 | 0.82 | .416 |

In sum, these results support the link between phonological awareness and reading performance and suggest that our two groups differ with regards to their reading ability and phonological awareness, but do not differ in important confounding factors such as vocabulary, age and hearing ability.

### 2.2 Decreased speech-brain coherence in subjects with developmental dyslexia

Subjects with (*N* = 18) and without DD (*N* = 26) listened to excerpts of an audiobook (“Das Märchen”, Goethe, 1795) narrated by a female speaker while magnetoencephalography (MEG) was recorded. Parts of the audiobook were noise-vocoded (see Figure 1A; 7-Chan, 3-Chan see *Methods – Stimuli* for details). At the end of each audio presentation, subjects were presented with two nouns from which they had to pick the one they perceived in the previous sentence (as in Chen et al., 2023; Hauswald et al., 2022; Schmidt et al., 2023). As expected task performance (i.e. hit rates; correct identification of the last spoken noun from two presented alternatives) significantly declined across all three intelligibility conditions for each group (*F*(1.46, 61.26) = 44.13, *p_ggeisser_* <.001, *η*^2^_P_ =.41). Aside from this main effect of intelligibility, there was no significant interaction between DD diagnosis and intelligibility (*F*(1.46, 61.26) = 1.11, *p_ggeisser_* =.335, *η*^2^_P_ =.02) and no significant difference between individuals with and without DD (*F*(1, 42) = 3.39, *p_ggeisser_* =.073, *η*^2^_P_ =.03). Bonferroni corrected pairwise comparisons for both individuals with DD and neurotypical controls revealed significant differences across all conditions (Original, 7-Chan & 3-Chan, all p<0.006; except for the difference between Original and 7-Chan (t(25)=1.28, p=.636 in neurotypical readers).

**Figure 1:**
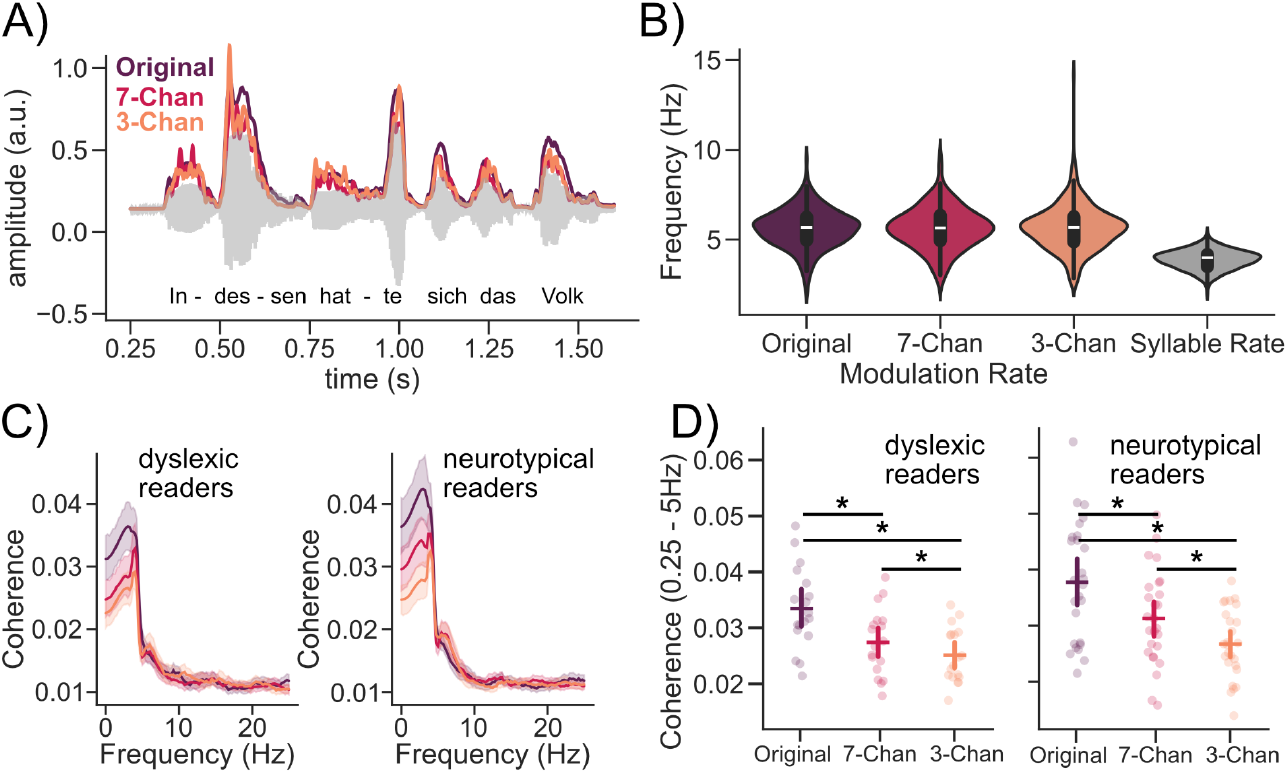
A) An exemplary snippet of Goethe’s “Das Märchen” with its corresponding original speech envelope and the envelopes of the same snippet, but vocoded at either 7 (less intelligible) or 3 channels (hardly intelligible). B) Syllable frequencies were extracted from original speech, while modulation frequencies were extracted from non-vocoded original speech segments, 7-channel-vocoded speech segments or 3-channel-vocoded speech segments respectively. C) Average speech-brain coherence spectra (in the 0-25 Hz range) with 95% confidence intervals for subjects without DD (n = 26) and subjects with a diagnosis of DD (n = 18) corresponding to the three intelligibility conditions: non-vocoded original speech (high intelligibility), 7-channel vocoded speech (moderate intelligibility) and 3-channel vocoded speech (low intelligibility). D) Coherence spectra from (C) averaged in the 0.25Hz - 5Hz range. Bars indicate 95% confidence intervals and asterisks denote significant Bonferroni corrected pairwise comparisons (*p*_bonferroni_ < 0.05).

To investigate how a loss of speech intelligibility via noise-vocoding influences the neural dynamics of speech tracking across both groups we measured the coherence between the speech envelope and the related cortical activity (see figure 1C). Considering the sharp drop in coherence after ∼5Hz we averaged coherence between 0.25 and 5Hz and tested for the effects of intelligibility and DD diagnosis, as well as their interaction, on neural speech tracking. This analysis revealed a significant main effect of intelligibility (*W*(1.61) = 38.36, *p* <.001, *η*^2^_P_ =.48). However, neither the interaction effect between DD diagnosis and intelligibility (*W*(1.61) = 0.37, *p* =.644, *η*^2^_P_ =.02) nor the main effect of DD diagnosis was significant (*W*(1) = 2.88, *p* =.089, *η*^2^_P_ =.07). Bonferroni corrected post-hoc pairwise comparisons revealed significant differences between all intelligibility conditions (*p*_bonferroni_ <.014). In sum, our results show that the decline of coherence with intelligibility did not differ significantly between the neurotypical and the dyslexic readers. Albeit not being statistically significant, subjects with DD still exhibited a trend towards reduced low-frequency (0.25 - 5 Hz) coherence when compared to subjects without DD (especially apparent in the Original & 7-Chan condition; see Figure 1C). Contrary to our predictions derived from previous studies, neither listening performance (Van Hirtum et al., 2021; Ziegler et al., 2009) nor the speech-brain coherence (Molinaro et al., 2016) of individuals with DD differed significantly from controls in difficult listening situations. However, coherence estimates without spectral parametrization may obscure the tracking of rhythmic information and thus fail to capture group differences (Schmidt et al. 2023).

### 2.3 Center Frequency of speech-brain coherences increases as intelligibility decreases in neurotypicals, but not in individuals with developmental dyslexia

To further investigate the differences in rhythmic low-frequency speech-brain coherence, we parametrized the coherence spectra using specparam (Donoghue et al. 2020). Afterwards we extracted the main peak in the low-frequency range (see *Methods – spectral parametrization*) and compared its “Peak Height”, “Center Frequency” and “Bandwidth” across the different levels of intelligibility and the group of subjects with DD and the matched controls. Those three parameters reflect complementary aspects of the neural tracking of rhythmic information, with peak height indexing the strength of coherence, center frequency indicating the temporal scale at which neural activity aligns to the speech envelope, and bandwidth capturing the range of frequencies contributing to this alignment (Schmidt et al., 2023).

For comparison of periodic speech tracking strength in the low frequency range between groups and intelligibility levels, we examined the “Peak Height” extracted via specparam. We found a significant main effect of intelligibility (*W*(1.91) = 11.24, *p* <.001, *η*^2^_P_ =.21). However, the interaction effect of DD diagnosis × intelligibility (*W*(1.91) = 0.52, *p* =.582, *η*^2^_P_ =.01) and the main effect of DD diagnosis (*W*(1) = 0.38, *p* =.538, *η*^2^_P_ <.01) were not significant. This shows that instead of being reduced in DD, the amplitude of periodic speech tracking in the low frequency peak in the speech-brain coherence spectra did not differ significantly between individuals with and without DD at any level of intelligibility (all *p*_bonferroni_ > 0.05). This result contradicts our previous expectations derived from the TSF (see https://osf.io/9f2xg; hypothesis 2), as we expected a reduction in the tracking of low-frequency rhythmic information in individuals with DD compared to controls.

For the Center Frequency (i.e. the temporal scale at which neural activity aligns to the speech envelope) we found a significant main effect of intelligibility (*W*(1.79) = 9.88, *p* <.001, *η*^2^_P_ =.27) and a significant interaction effect between DD diagnosis and intelligibility (*W*(1.79) = 3.72, *p* =.029, *η*^2^_P_ =.13). Again, the main effect of DD diagnosis was not significant (*W*(1.00) = 2.98, *p* =.084, *η*^2^_P_ =.11) suggesting no overall difference in center frequency between individuals with and without DD. Bonferroni corrected nonparametric post hoc pairwise comparisons are summarised in Figure 2B, indicating that in neurotypical subjects center frequency increases as intelligibility decreases. This difference in center frequency was not significant in subjects with DD. To further hone into the shift in center frequency we fitted linear regression models separately for each subject and extracted the slopes of these individual linear models across all three intelligibility conditions. These slopes indicating “shifts” in center frequency across vocoding conditions were then compared between subjects without DD and those with a DD diagnosis in a Wilcoxon Rank Sum test. As expected, individuals without DD (Δ Slope*_Mdn_* = 0.32 Hz) showed a bigger shift in the temporal scale of the neural tracking than those with DD (Δ Slope*_Mdn_* = 0.07 Hz): *W* = 339, *Z* = 2.51, *p* =.013, *r* =.35. This result aligns with our expectations that the frequency of the neural tracking does not adapt to changes in intelligibility (hypothesis 1). However, contrary to our expectation we find that the temporal scale of this tracking is aligned closer to the syllable rate than to the modulation rate in DD across all vocoding levels (see Figure 1B & 2B; hypothesis 1).

**Figure 2:**
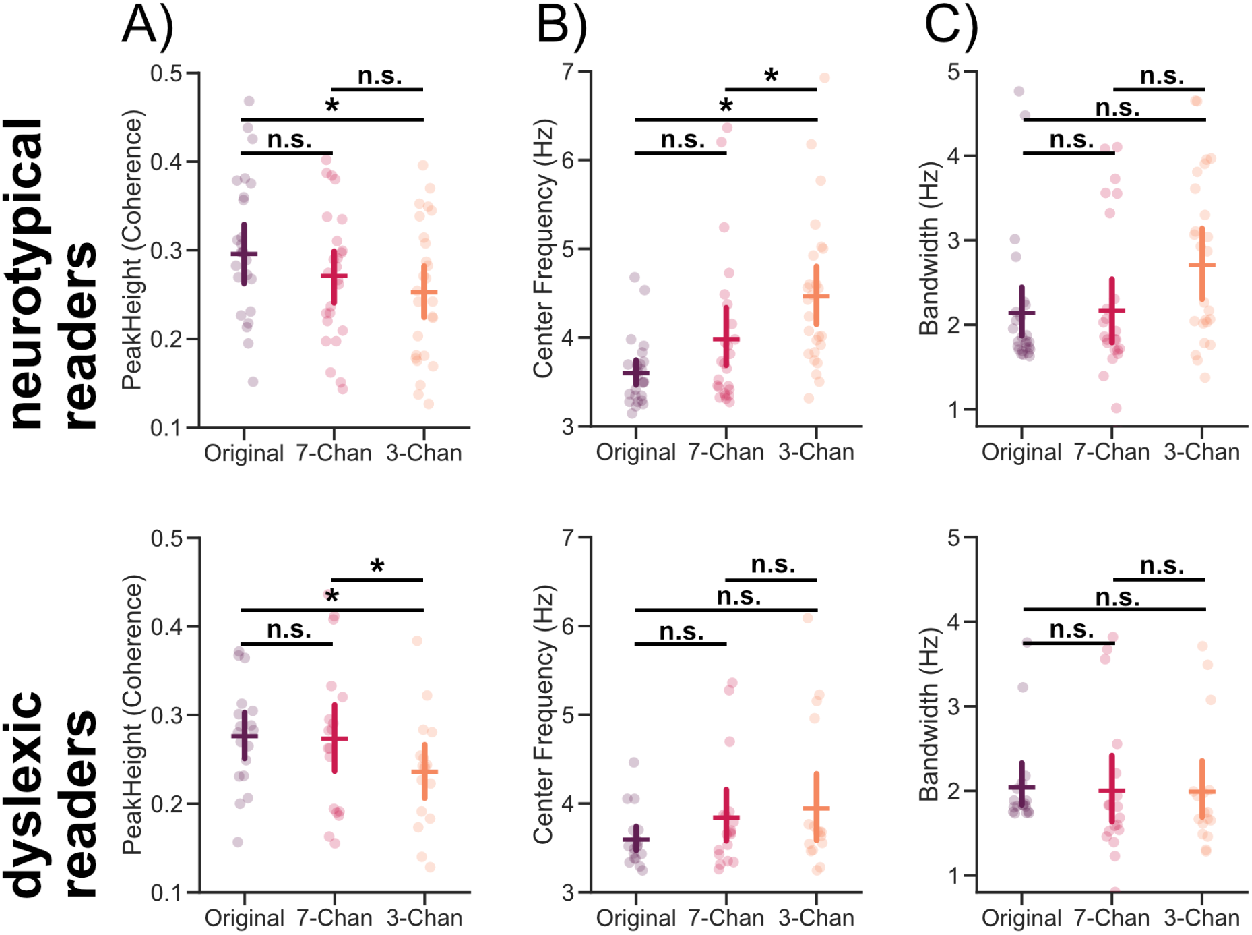
Periodic parameters extracted from the coherence spectra for the two groups and three conditions. A) Peak height of coherence B) Center frequency and C) bandwidth of main peak. A-C) Each dot represents one of the 42 subjects (as two participants lacked an identifiable peak in the low frequency range: *N_no DD_*=25, *N_DD_*=17) whose grand average coherence spectra had a low frequency peak clearly identifiable via the specparam algorithm in all three intelligibility conditions. Bars indicate 95% confidence intervals and asterisks denote significant Bonferroni corrected pairwise comparisons (*p*_bonferroni_ < 0.05).

Additionally, we compared the bandwidth values of the main low-frequency speech-brain coherence peaks. It revealed a significant interaction effect between DD diagnosis and intelligibility (*W*(1.89) = 3.58, *p* =.030, *η*^2^_P_ =.12), while neither the main effects of intelligibility (*W*(1.89) = 1.01, *p* =.359, *η*^2^_P_ =.09) nor those of DD diagnosis (*W*(1.00) = 3.23, *p* =.072, *η*^2^_P_ =.14) were significant. Bonferroni corrected nonparametric post hoc pairwise comparisons are summarised in Figure 2C, showing that bandwidth did not differ across the three vocoding conditions both for neurotypical and dyslectic readers.

In sum, these findings show that the center frequency of speech–brain coherence robustly increases as intelligibility decreases in neurotypical readers—a pattern consistently observed in previous work (Schmidt et al., 2023; Chen et al., 2023)—but not in individuals with DD. Yet, it remains unclear if these neural changes also correspond to any behaviorally measurable DD-related deficits?

### 2.4 Center Frequency shifts emerge as the sole neural predictor of reading and phonological awareness

To investigate, if any of the (parametrized) changes in low-frequency neural tracking correspond to DD-related deficits, we correlated the different parameters extracted from the coherence spectrum with indices of reading performance and phonological awareness.

To better capture the change across the different parameters reflecting low-frequency coherence we first fit linear regression models per parameter (e.g. as mentioned above for center frequency) separately for each subject across all three intelligibility conditions and extracted the slopes of these individual linear model fits. These slopes reflect the change in a coherence parameter across the different levels of intelligibility. Afterwards, Shepherd’s *pi* correlations (i.e. spearman’s rank-order correlations after accounting for the effect of bivariate outliers) between the slopes and the cognitive tests for reading ability and phonological awareness were calculated (see Figure 3AB).

**Figure 3:**
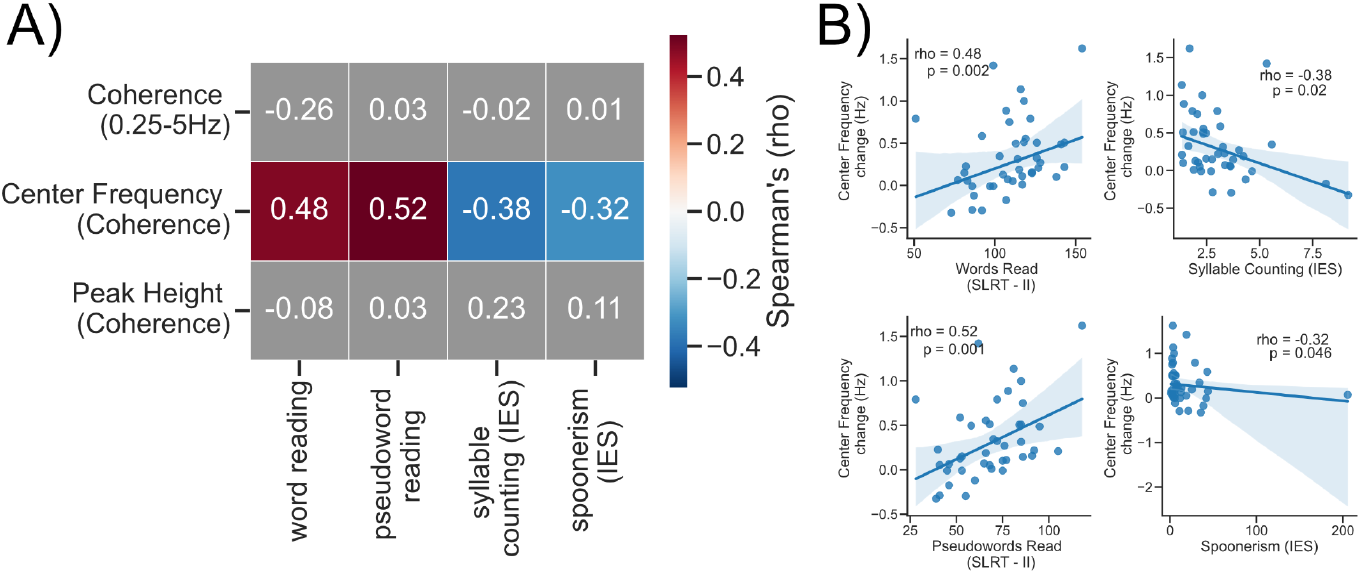
*Center-frequency shifts are associated with reading performance and phonological awareness.* A) Shepherd’s *pi* correlations were calculated between coherence measures (averaged Coherence (0.25-5Hz); center frequency; peak height) and behavioral measures of reading ability and phonological awareness. Among the coherence measures, only center frequency showed reliable associations with behavior. Participants who exhibited stronger shifts in the center frequency of speech-brain coherence across vocoding levels read (pseudo)words more quickly and were more efficient in successfully performing phonological awareness tasks. B) Correlations between center frequency and behavioral measures. Shepherd’s *pi* correlations were used to reduce the influence of outliers (e.g. spoonerism scores in panel B).

The results of this analysis show that task-related shifts in the center frequency of speech-brain coherence are associated with the performance in reading and phonological awareness tasks. Subjects with a faster word and pseudoword reading speed also exhibited stronger shift in center frequency with increased vocoding (*r_word’s read_*=0.48, *p _word’s read_*=0.002 & *r_pseudoword’s read_*=0.52, *p_pseudoword’s read_*=0.001). Furthermore, the inverse efficiency scores of syllable counting and spoonerism (phonological awareness tasks) were negatively related to shift magnitude (*r_Syllable counting_*=-0.38, *p _Syllable counting_*=0.02 & *r_spoonerism’s_*=-0.32, *p_spoonerism’s_*=0.046). This shows that individuals with a stronger shift in center frequency magnitude were more efficient in successfully performing “spoonerisms” and “syllable counting”. All the other parameters – that also showed some significant differences across intelligibility levels (see Figure 1 & 2) – were not significantly related to reading or phonological awareness (see Figure 3A).

In sum, these results show that subjects that exhibited a stronger shift in the center frequency of their speech-brain coherence across different vocoding levels were able to read more (pseudo-)words in a shorter time and were more efficient in successfully performing phonological awareness tasks. Importantly, neither age (*r*=.17, *p*=.25), hearing ability (*r*=.18, *p*=.25) nor vocabulary (*r*=-.01, *p*=.96) were significantly related to the shift’s magnitude. This result mostly aligns with our expectations to find the magnitude of the shifting effect to be related to performance in reading and phonological awareness tasks (see preregistration exploratory analysis 1).

## 3. Discussion

This study set out to investigate whether individuals with developmental dyslexia (DD) differ from matched controls in their neural tracking of slow (<10 Hz) speech rhythms under varying levels of intelligibility. Behaviorally, we confirmed robust group differences: individuals with DD showed marked impairments in reading performance and phonological awareness, consistent with prior evidence that such difficulties persist into adulthood (Bruck, 1992; Reis et al., 2020). Importantly, the two groups did not differ in vocabulary, age, or hearing ability, indicating that the observed difficulties in reading and phonological awareness cannot be attributed to general knowledge or other sensoric deficits (e.g. hearing loss). Using noise-vocoded speech and speech–brain coherence analysis, we isolated the main peak of the low-frequency coherence spectrum (<10 Hz; Schmidt et al., 2023). Interestingly, we found that not a reduction in the overall magnitude of this peak, but rather reduced flexibility in its center frequency across vocoding levels, was the only neural-tracking measure linked to phonological awareness and reading ability. This partly contradicts predictions derived from the TSF: instead of a general reduction in the tracking of rhythmic speech features, DD appears to be characterized by an inflexibility in adjusting neural rhythms as speech intelligibility decreases.

### 3.1 From amplitude differences to center-frequency shifts

A core prediction of the TSF is that in DD neural oscillations in the low frequency range (<10Hz; namely in the delta and theta band) fail to properly entrain to corresponding rhythmic information in the speech envelope. This prediction is typically operationalized as a reduction in coherence within these frequency bands (Abrams et al., 2009; Molinaro et al., 2016; Goswami, 2011, 2018). However, just as non-zero power in a spectrum does not necessarily indicate the presence of a genuine oscillation (Donoghue et al. 2020), a non-zero coherence value does not in itself guarantee the presence of coupling between two oscillators (Srinath & Ray, 2014; Schmidt et al., 2023).

We therefore parametrized the coherence spectra, extracted the main peak in the low-frequency range (<10Hz) and compared its “Peak Height”, “Center Frequency” and “Bandwidth” across the different levels of intelligibility and the group of subjects with DD and the matched controls. Those three parameters reflect complementary aspects of the neural tracking of rhythmic information, with peak height indexing the strength of coherence, center frequency indicating the temporal scale at which neural activity aligns to the speech envelope, and bandwidth capturing the range of frequencies contributing to this alignment (Schmidt et al., 2023).

Contrary to our prediction of reduced “Peak Height” in individuals with DD (hypothesis 2; https://osf.io/9f2xg), we instead found that the center frequency of speech-brain coherence increases with decreasing intelligibility in neurotypical readers – a robust pattern reported previously (Schmidt et al., 2023; Chen et al., 2023) – but not in readers with DD. This suggests that the tracking of rhythmic fluctuations in speech is not simply reduced by decreasing signal quality, but that the brain of neurotypical readers flexibly prioritizes different temporal cues in the acoustic signal depending on how intelligible the speech is (Schmidt et al., 2023). This process is largely absent in individuals with DD. Importantly, without parametrization, such increases in center frequency can sometimes appear as reductions in coherence magnitude in a specific band e.g. delta or theta. This raises the possibility that what has often been described as reduced neural tracking – in work building on the TSF – may in fact reflect a lack of flexibility in adapting neural sampling to changing acoustic cues (Goswami, 2011, 2018, 2019).

### 3.2 Center-Frequency Flexibility and Its Behavioral Relevance

In line with our preregistered hypothesis (see https://osf.io/9f2xg), the intelligibility-dependent center-frequency shift was significantly reduced in individuals with DD. This finding supports the TSF insofar as oscillatory alignment in the low frequency range is atypical in DD (see Goswami, 2011). However, unlike a general reduction in tracking strength, the deficit we observed was characterized by reduced flexibility: neural tracking in DD remained more rigidly aligned to the syllable rate, rather than dynamically adjusting to faster envelope fluctuations under degraded listening conditions. Notably, this pattern runs counter to our initial expectation that individuals with DD would show a rigid alignment to the modulation rate instead, reflecting a bias toward purely acoustic rather than linguistic cues. This rigidity in low frequency synchronization is consistent with recent findings. Zhang et al. (2022), for example, showed that individuals with DD exhibit increased integration in theta-band connectivity during word tracking, suggesting an overly synchronized and less adaptable theta network. Taken together, these results indicate that DD is not defined by a reduction of low-frequency neural speech tracking, but rather by tracking that is too inflexible, preventing an adaptive alignment to changing acoustic cues.

### 3.3 Behavioral Correlates

A key strength of our study is the link between neural tracking flexibility and behavior. While amplitude-based measures of coherence have often been assumed to underlie reading performance, we found no consistent associations between reading-related skills and broadband or peak amplitude changes across intelligibility conditions. By contrast, the magnitude of the low frequency center-frequency shift was reliably related to both reading and phonological awareness: subjects that exhibited a stronger shift in the center frequency of their speech-brain coherence across different vocoding levels were able to read more (pseudo)words in a shorter time and were more efficient in successfully performing phonological awareness tasks. These findings suggest that it is not the strength of neural tracking per se, but the flexibility of periodic sampling that is most critical for reading-related skills. More specifically, changes in center frequency across conditions, rather than changes in amplitude height, were most predictive of individual performance.

### 3.4 Interpretation within the TSF

Our findings refine the TSF in important ways. The original framework emphasizes impaired entrainment of neural oscillations in the delta–theta range as a core feature of DD (Goswami, 2011, 2018). Consistent with this, we found evidence for reduced adaptability of low-frequency (theta-range) tracking. However, when coherence spectra were parameterized, no group differences emerged in peak amplitude. This indicates that the key deficit lies not in oscillatory gain (how strongly neural activity couples to speech in a specific band), but in sampling flexibility (which information in the signal is sampled by neural activity). Because coherence can conflate periodic and aperiodic contributions, amplitude-based measures may obscure this distinction (Srinath & Ray, 2014, Schmidt et al. 2023). By isolating periodic peaks in the spectrum, we show that flexibility in the center-frequency of a peak provides a cleaner index of adaptive sampling. This may also explain why prior studies have reported mixed results: delta-band differences appear more robust, whereas theta-band amplitude effects are inconsistent. Our center-frequency shift result therefore points to altered adaptability of periodic tracking (rather than reduced periodic coupling per se), dovetailing with TSF and helping to reconcile mixed theta findings. Moreover, the TSF has so far focused primarily on rhythmic amplitude modulations without explicitly distinguishing periodic from aperiodic processes. If these reflect distinct neural mechanisms, they may contribute in complementary yet dissociable ways to speech encoding. Future refinements of the TSF should incorporate this distinction to more fully capture the complexity of speech–brain coupling in developmental disorders.

### 3.5 Limitations

Several limitations of the present study should be noted. First, our analyses were based on coherence spectra averaged across channels. While this approach provides a global measure of neural tracking, more fine-grained effects might emerge when investigating channel-specific patterns, both in the unparameterized and parameterized data. Future work should therefore assess whether different coherence differences emerge in a spatially resolved manner and how they contribute to the grand averaged group effects observed herein. Second, our evidence is correlational and limited to an adult sample. Although we demonstrate that reduced flexibility in center-frequency shifts is associated with phonological awareness and reading performance, we cannot infer causality from these relationships. It remains to be determined how this neural marker develops over time and whether it can serve as an early predictor of dyslexia in children. Longitudinal and developmental studies will be essential to establish the prognostic value of center-frequency flexibility to varying acoustic inputs.

## 4. Conclusion

Taken together, our findings add to the growing body of evidence that individuals with DD exhibit atypical neural tracking to speech rhythms below 10 Hz. Contrary to predictions of a general reduction in neural tracking, we observed preserved strength in neural tracking but reduced flexibility: individuals with DD showed rigid sampling of low-frequency speech components, remaining closely aligned to the syllable rate even when intelligibility decreased. This suggests that the critical impairment is not the overall ability to track rhythmic features of speech, but the adaptability of neural tracking to changing acoustic demands. By identifying center-frequency shifts as the spectral parameter most strongly associated with reading and phonological awareness, our study calls for a refinement of the TSF and highlights temporal flexibility as a potential neural marker of dyslexia-related deficits.

## 5. Methods

The initial plan for acquisition and analysis was pre-registered before starting data collection and can be accessed via the following link: https://osf.io/9f2xg.

### 5.1. Participants

A total of 44 participants between the ages 18 and 50 years completed the behavioral assessment and the MEG recordings. In the DD group there were 18 individuals who had previously been diagnosed with DD (female: 10, male: 7, other: 1) and in the control group were 26 individuals without such a diagnosis (female: 14, male: 12, other: 0). The average age for participants with a diagnosis was 24.94 years (*SD* = 6.75) and for those without one it was 26.88 years (*SD* = 8.89) which did not differ between groups *t*(41.58) = 0.82, *p* =.415, *d* = 0.24. One participant with DD and one without DD had to be excluded from tests on parameters extracted using specparam due to a missing low frequency peak in one condition. All participants were required to be in good health. Current physical (e.g. cold), psychiatric (e.g. depression, substance use disorder, schizophrenia) and neurological (e.g. history of stroke, epilepsy, dementia) conditions led to exclusion from the study. All participants provided informed consent and were compensated monetarily with 10€ per hour spent at the lab. Though most participants were university students, none of them asked for compensation via course credits. Participation was voluntary and in accordance with the declaration of Helsinki as well as with the statutes of the University of Salzburg. The study procedure was approved by the ethical committee of the University of Salzburg (GZ 22/2016).

### 5.2. Stimuli

For stimulation during the MEG recordings we reused 24 audio files of a female speaker reading the original German version of Goethe’s “The Green Snake and the Beautiful Lily” (“Das Märchen”, 1795). These recordings were created as continuous speech stimuli for previous studies on speech tracking (Hauswald et al., 2022; Schmidt et al., 2023; Chen et al. 2023). By noise-vocoding of some stimuli, we realised 3 levels of intelligibility: Unaltered and clearly intelligible non-vocoded original speech (original), less intelligible 7-channel vocoded speech (7-Chan) and hardly intelligible 3-channel vocoded speech (3-Chan). Fewer vocoding channels result in a more rudimentary speech signal, which is less intelligible.

Approximately 2/4s of audio runtime were vocoded utilising the vocoder toolbox for MATLAB (Gaudrain, 2016) as described by Hauswald et al. (2022):

“For the vocoding, the waveform of each audio stimulus was passed through two Butterworth analysis filters (for 7 and 3 channels) with a range of 200–7000 Hz, representing equal distances along the basilar membrane. Amplitude envelope extraction was done with half-wave rectification and low-pass filtering at 250 Hz. The envelopes were then normalized in each channel and multiplied with the carrier. Then, they were filtered in the band and the RMS (Root Mean Square) of the resulting signal was adjusted to that of the original signal filtered in that same band. Auditory stimuli were presented binaurally using MEG-compatible pneumatic in-ear headphones (SOUNDPixx, VPixx technologies).”

The sequence of all presented vocoded and non-vocoded audiobook segments was randomized individually for each subject, resulting in a variety of sequences that did not follow the fairy tale’s original storyline. To ensure similar duration of stimulation at each intelligibility level, the assignment of audiobook segments to vocoding options, was controlled at subject level by boundary conditions for total audiobook durations in each intelligibility condition.

Participants were instructed to listen attentively to the presented continuous speech stimuli. Each one of the 24 continuous speech stimuli (presented in 8 blocks à 3 stimuli) ended with a four-word sequence that contained a two-syllable noun. At the end of each stimulus, participants were presented with 2 written two-syllable nouns via a screen and were asked to identify the word they had (most likely) heard last. This was done to keep participants’ attention on the stimuli and to facilitate comparison of intelligibility between conditions. To prevent participants from anticipating the end of continuous speech stimuli and adapting levels of attentiveness accordingly, stimulus durations ranged from 16 s to 3 min 14 s.

### 5.3. Data Acquisition

Participants were required to attend the lab twice within 4 weeks. Once for a 1.5-hour cognitive screening and once for the 2.5-hour collection of MEG data during audiobook presentation. The order of those sessions was flexible depending on both the researcher’s and the participant’s schedules. A study assistant screened volunteers for DD diagnoses and selected age matched controls. To ensure blinding, the investigator received only contact information and was not informed of participants’ diagnostic status (neurotypical vs. dyslectic readers).

#### 5.3.1. Cognitive Tasks – Session

During the cognitive screening participants completed 5 test and tasks in the very order they are described here: To enable controlling for potentially confounding top-down influences of vocabulary on speech tracking acuity, the Wechsler Adult Intelligence Scale IV’s vocabulary test (WAIS-IV: Wortschatztest, Pearson GmbH) was administered first (Wechsler, 2012). Next, participants completed the 2 short reading tasks contained in the Salzburger Lese- und Rechtschreibtest (SLRT-II: Lesetest, Hogrefe AG): the word-reading task and the pseudoword-reading task (Moll & Landerl, 2014). In both these tasks participants were given 1 minute to read as many words or pseudowords as they could. At last, phonemic and phonological awareness were assessed via 2 self-developed tasks: These tasks were inspired by child the oriented tests BAKO (Stock et al., 2017) and QUIL-D (Schnitzler, 2015) and adapted for adults by choosing less frequent real words (spoonerisms) or inventing new pseudowords (syllable counting). Participants completed a spoonerisms-task with 27 items in which they had to swap the first phonemes of two words (e.g. “Zahl Schuh” → “Schal zu”) nd a syllable counting-task with 20 items in which they counted syllables of newly invented German pseudowords (e.g. “Wostimetikon” → 5 syllables). The former task requires the ability to identify and manipulate individual sounds in spoken words and mature capacities of verbal working memory, while the latter task assesses more general phonological awareness (de Jong & van der Leij, 2003; Smail et al., 2021), which comes first in the general trajectory of phonological awareness development (Anthony & Francis, 2005). In both the spoonerism-task and the syllable counting-task, response times were recorded via a custom MATLAB-application (MathWorks, 2020). For this purpose, participants were required to press a button immediately before answering. Answers were evaluated for correctness on site by the instructor, also via button press.

#### 5.3.2. MEG – Session

At the start participants first completed an online hearing assessment (https://salzburghoert.sbg.ac.at/). They then underwent pure-tone audiometry (AS608 Screening-Audiometer, Interacoustics) and preparation for MEG recordings, which involved attaching 5 coils to track head position, 4 electrodes measuring vertical and horizontal eye movements, 2 electrodes measuring heart activity and 1 ground electrode. Before entering the magnetically shielded room (AK3B, Vakuumschmelze), the head shape of each participant was acquired with at least 300 digitized points on the scalp, including fiducials (nasion, as well as left and right preauricular points) by a FASTRAK system (Polhemus). Magnetic brain activity was recorded via a 306-channels MEG system, 204 planar gradiometers and 102 magnetometers (Elekta-Neuromag^®^ TRIUX, Elekta Oy) with a sampling rate of 1 kHz while participants were listening to a total of 24 vocoded and non-vocoded continuous speech stimuli.

### 5.4. Data Analysis

Data analysis steps were performed similarly as in prior studies on speech tracking that were conducted by Schmidt et al. (2023) and Chen et al. (2023).

#### 5.4.1. Extracting Speech Envelopes

Acoustic speech envelopes were extracted from all 24x3 continuous speech stimuli, at their actual intelligibility levels (Original, 7-Chan and 3-Chan), using the Chimera toolbox in MATLAB (http://research.meei.harvard.edu/chimera/More.html) by Bertrand Delgutte (Smith et al., 2002). The “filter-Hilbert-method” (Cohen, 2014; Ding et al., 2017) was then applied. For this purpose, 9 frequency bands in the range of 100 Hz to 10 kHz were constructed. These frequency bands were selected in a manner equidistant on the basilar membrane, which exhibits almost logarithmic frequency spacing properties, an organisational pattern that is known as the “cochlear map” (Gross et al., 2013; Li et al., 2021; Smith et al., 2002). Each continuous speech stimulus was band-pass filtered twice in these 9 frequency bands (once forward and once in reverse), using 4^th^-order Butterworth filters. Subsequently, envelopes were calculated for each of the 9 band-pass filtered continuous speech signals, as absolute values of the Hilbert Transform (Hilbert, 1912): abs(hilbert(speech_signal)) in MATLAB (MathWorks, 2020). The resulting 9 narrow-band envelopes were averaged to generate one broadband envelope and downsampled to 200Hz. As expected, the resulting speech envelopes were similar across the three intelligibility-conditions (Figure 1A). Speech envelopes were later used to estimate acoustic modulation rates and calculate speech-brain coherence.

#### 5.4.2. Estimating Modulation Rates and Syllable Rates

Estimates of the acoustic modulation rate were calculated for all 24 continuous speech stimuli (Original, 7-Chan and 3-Chan).We used 6 second envelope segments to compute power spectra of the broadband envelope using Welch’s method (Welch, 1967). Power spectra were then parametrized in their aperiodic and periodic components using the specparam algorithm (Donoghue et al. 2020; with the following settings). The center frequency of the peak with the highest power in the low frequency range (0.25-16Hz) for each 6 second segment was extracted (Schmidt et al., 2023), which is henceforth referred to as “acoustic modulation rate”. Acoustic modulation rates were averaged individually for each participant over all corresponding 6 second segments for each of the 3 intelligibility-conditions (Figure 1B). Grand average modulations rates did not differ across the 3 intelligibility-conditions (*F*(2, 1098) = 0.03, *p* = 0.967, *η*^2^ = 0.00). The syllable rates were computed individually for all 24 clearly intelligible non-vocoded speech stimuli using Praat (de Jong & Wempe, 2009) and found to vary between 3.99 and 4.60 Hz with a median of 4.21Hz (https://osf.io/w3r7t/).

#### 5.4.3. Preprocessing of MEG Data

The raw MEG data was Maxwell filtered using a SSS (Signal Space Separation) algorithm (Taulu & Simola, 2006) implemented in the MaxFilter^TM^ software (version 2.2.15) provided by the manufacturer (Electa Oy) of our MEG system to remove external magnetic interference from the MEG signal. Maxwell filtering also realigned data to a common standard head position (with default MaxFilter^TM^ parameters). The filtered data were then further analysed using the FieldTrip toolbox (Oostenveld et al., 2011) and custom-built MATLAB (MathWorks, 2020) routines: The MEG data was low-pass filtered at 40Hz and high-pass filtered at 0.1 Hz using FIR (finite impulse response) filters (Kaiser windows) to eliminate muscle artefacts as well as scanner drifts. Physiological artifacts (eye movements and cardiac activity) were extracted from the data by calculating 50 independent components via ICA, using the runica method implemented in the FieldTrip toolbox (Oostenveld et al., 2011). The 50 components were visually inspected. Those that consistently showed eye movements or heartbeats were removed from the data. ICA was completed separately for each of the 8 blocks available per subject. On average 3.71 components were removed per subject (SD = 1.05). Next, we only selected magnetometers for further analyses and downsampled the data to 200 Hz to reduce computational load, since we were interested in primarily low frequency activity. Then, speech envelopes of the corresponding intelligibility condition were appended to the MEG data structure, while accounting for stimulation delay relative to triggers of 16ms (measured during piloting with The Black Box Toolkit v2 (Richard, 2003)). Aligned data matrices were cut into segments of 4 second duration. The number of segments per intelligibility condition was equalized across intelligibility conditions (Original, 7-Chan and 3-Chan) to the lowest number of 4 second segments available per subject. This resulted in some variability in the number of audiobook segments (available for each intelligibility condition) across subjects ranging from 170 to 231 segments (*M* = 209.76, *SD* = 12.64).

#### 5.4.4. Speech-Brain Coherence

We calculated multi-taper frequency transformations on each of the epochs, using the following parameters: dpss taper sequence 0–25 Hz in 0.25 Hz steps, 4 Hz smoothing, no baseline correction. The resulting power spectra of the MEG signal and the speech envelope at their specific intelligibility levels were then used for calculation of the speech-brain coherence at each of the 102 magnetometers individually:

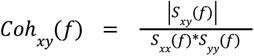

where Sxy is the cross spectral density between the MEG signal and the speech envelope, while Sxx and Syy are auto spectral densities of the MEG signal and the speech envelope respectively. For coherence calculation between the MEG signal at each virtual sensor and the acoustic speech envelope, matching frequency bins of 0.25 Hz were chosen.

#### 5.4.5. spectral parametrizations

We extracted the most prominent low frequency peaks from coherence spectra using the specparam-algorithm (https://fooof-tools.github.io/fooof/ Donoghue et al., 2020). Peak extraction was performed on the coherence spectra averaged across all 102 magnetometers. Averaged coherence spectra were flattened by subtracting an aperiodic broadband component. Gaussian model fits were computed to extract the most prominent peak in the range between 1 and 10.5 Hz from each channel types mean coherence spectra. Only peaks exceeding a threshold of 2 standard deviations (default setting) relative to the aperiodic slope (peak_threshold = 2.0) were included. For some average coherence spectra no peak could be identified (*N* = 2). For each subject, peak height, bandwidth, and center frequency of the peaks were extracted three times (once per intelligibility condition).

#### 5.4.6. Statistical Analysis

All statistical analyses were completed in R using one or more of the following packages: base (R Core Team, 2024), stats (R Core Team, 2024), BayesFactor (Morey & Rouder, 2018), coin (Hothorn et al., 2008), lme4 (Bates et al., 2015), rstatix (Kassambara, 2023), nparLD (Noguchi et al., 2012). All figures were created in Python using seaborn (Waskom, 2021) and matplotlib (Hunter, 2007).

##### 5.4.6.1. Cognitive Tasks

Scores on all five cognitive tasks (vocabulary, word reading, pseudoword-reading, spoonerisms and syllable-counting), age and pure tone audiometry outcomes were compared between subjects with and without DD via independent samples Welch’s t-tests (package: stats), which are robust against heteroskedasticity and slight violations of normality, as long as there is no cumulation of multiple violations and sample sizes are approximately equal (Ahad & Yahaya, 2014; Curtis, 2024). For the spoonerisms as well as the syllable counting task, we combined accuracy (number of correctly solved items) and response times into a single inverse efficiency score (IES) per task (Townsend & Ashby, 1983). IES were calculated for each participant by dividing the mean response time for correct solutions by the proportion of correctly solved items.

##### 5.4.6.2. Behavioral Task

Hit rates for the three intelligibility conditions were compared within groups and between groups via a two-way mixed ANOVA and Bonferroni corrected t-tests (package: rstatix). Due to aiming for overall similar stimulus duration of the three conditions, the absolute number of trials per condition differed (mainly assignment of the shorter trials, i.e. less than three minutes). Statistical comparisons therefore were only calculated for the twelve long (>three minutes) trials that always were assigned equally to one of the three conditions (four per condition). Bayes factors, comparing individuals with DD to controls, were calculated using the “ttestBF” function (package: BayesFactor) with the same default medium Cauchy prior (r = 0.707).

##### 5.4.6.3. Speech-Brain Coherence

We investigated speech tracking in participants with and without DD, with a set of 4 nonparametric two-way mixed ANOVAs using the Wald-type test (as the statistical assumptions for parametric ANOVAs were not met for all coherence related analyses; package: nparLD). One group variable (DD and no DD) and one repeated-measures variable (Original, 7-Chan and 3-Chan) were used to compare the averaged low frequency coherence (0.25-5Hz) as well as the peak parameters (center frequency, peak height and bandwidth) of the main low frequency peak extracted from the coherence spectra via specparam to examine the impact of DD diagnosis and intelligibility.

##### 5.4.6.4. Cognitive Tasks X Speech-Brain Coherence

Ultimately, we analysed the association between the number of correctly read words and pseudowords (measure to assess DD) and speech-brain coherence measures (average low-frequency coherence, peak-height and center frequency shift). For the coherence measures, slopes across all three vocoding conditions were calculated within-subject using linear models; afterwards Spearman’s rank correlation coefficients (*r*_s_) between the Cognitive Tasks and these within-subjects changes of speech-brain coherence were calculated.

